# Routing of non-mnemonic hippocampal ripples to subcortical structures

**DOI:** 10.1101/610402

**Authors:** David Tingley, György Buzsáki

## Abstract

The mnemonic functions of hippocampal sharp wave ripples (SPW-Rs) have been studied extensively. Because hippocampal outputs affect not only downsteam cortical structures but also subcortical targets, we examined the impact of SPW-Rs on the firing patterns of lateral septal (LS) neurons in behaving rats. A large fraction of SPW-Rs were temporally locked to high frequency oscillations (HFOs; 120-180 Hz) in LS. SPW-R to LS HFOs coupling was strongest during NREM sleep, followed by waking immobility. However, coherence and spike-LFP coupling between the two events were low, suggesting that HFOs are generated locally within the LS GABAergic population. This hypothesis was supported by optogenetic induction of HFOs in LS. Spiking of LS neurons was independent from the sequential order of spiking in SPW-Rs but, instead, correlated strongly with the magnitude of excitatory synchrony of the hippocampal output. Thus, LS is strongly activated by SPW-Rs and may convey hippocampal population events to its motivation and action-inducing targets in the hypothalamus and brainstem.

## Introduction

Current theories on the function of hippocampal sharp-wave ripples (SPW-Rs) relate mainly to the consolidation of recent experiences and planning of future actions (Buzsáki, 2015; Foster, 2017a; Jadhav et al., 2009; Pfeiffer, 2018). Specifically, sequential firing patterns within SPW-Rs are thought to ‘replay’ recently experienced episodes and planning of routes (Diba and Buzsáki, 2007; Gupta et al., 2010; Karlsson and Frank, 2009; Pfeiffer and Foster, 2015; Skaggs and Mcnaughton, 1996). Both of these ideas tacitly assume that the main function of the SPW-R is to convey content-specific hippocampal messages to the neocortex (Ji and Wilson, 2016; Ólafsdóttir et al., 2016; Peyrache et al., 2009; Rothschild et al., 2017; Sirota et al., 2003a; Wang and Ikemoto, 2016) or subcortical structures (Dragoi et al., 1999; Girardeau et al., 2017; Gomperts et al., 2015; Lansink et al., 2009; Pennartz, 2004; Sjulson et al., 2018). While it is clear that hippocampal SPW-Rs co-occur with brain wide fluctuations in activity (Logothetis et al., 2012), many of these regions do not have strong monosynaptic connections with the hippocampal formation. We reasoned that such coupling relationships likely emerge through multi-synaptic brain circuits and sought to examine which brain regions may serve as a conduit, receiving direct hippocampal inputs, thus allowing for SPW-R propagation out of the hippocampus.

A major target of the hippocampal system is the lateral septum (LS), where the density of hippocampal afferents is ∼20-fold higher when compared to entorhinal regions or ∼180-fold higher when compared to prefrontal targets (Tingley and Buzsáki, 2018). This makes the LS the recipient of massive convergent excitation from the hippocampal-subicular-entorhinal population. This major anatomical connection may convey hippocampal activity to its hypothalamic, mesencephalic and brainstem projections (Swanson and Cowan, 1979). In light of these anatomical connections, we set out to examine the hippocampal-LS circuit by simultaneously recording activity patterns of the hippocampus and LS in the behaving rat.

## Results

Using silicon probes, we simultaneously recorded neurons in the hippocampus (CA1 and CA3) and LS while animals slept and performed behavioral tasks. Details of the surgical procedures and behavioral experiments have been published (Buzsáki and Tingley, 2018). During initial recordings in LS (N = 6 rats), we observed transient high frequency oscillations (HFOs) in the 120-180 Hz frequency range in each animal (Figure 1A, B) and an associated robust increase in the firing probability of LS neurons (Figure 1C). On average, 16% of LS neurons participated in any given event, firing an average of 0.23 spikes per event (Figure S1A, B). The vast majority of LS neurons (1840 of 1989; 92%) were significantly phase-locked to the 120-200 Hz frequency band, with a strong unimodal phase preference for the trough of each cycle (Figure 1D,E).

**Figure 1.**
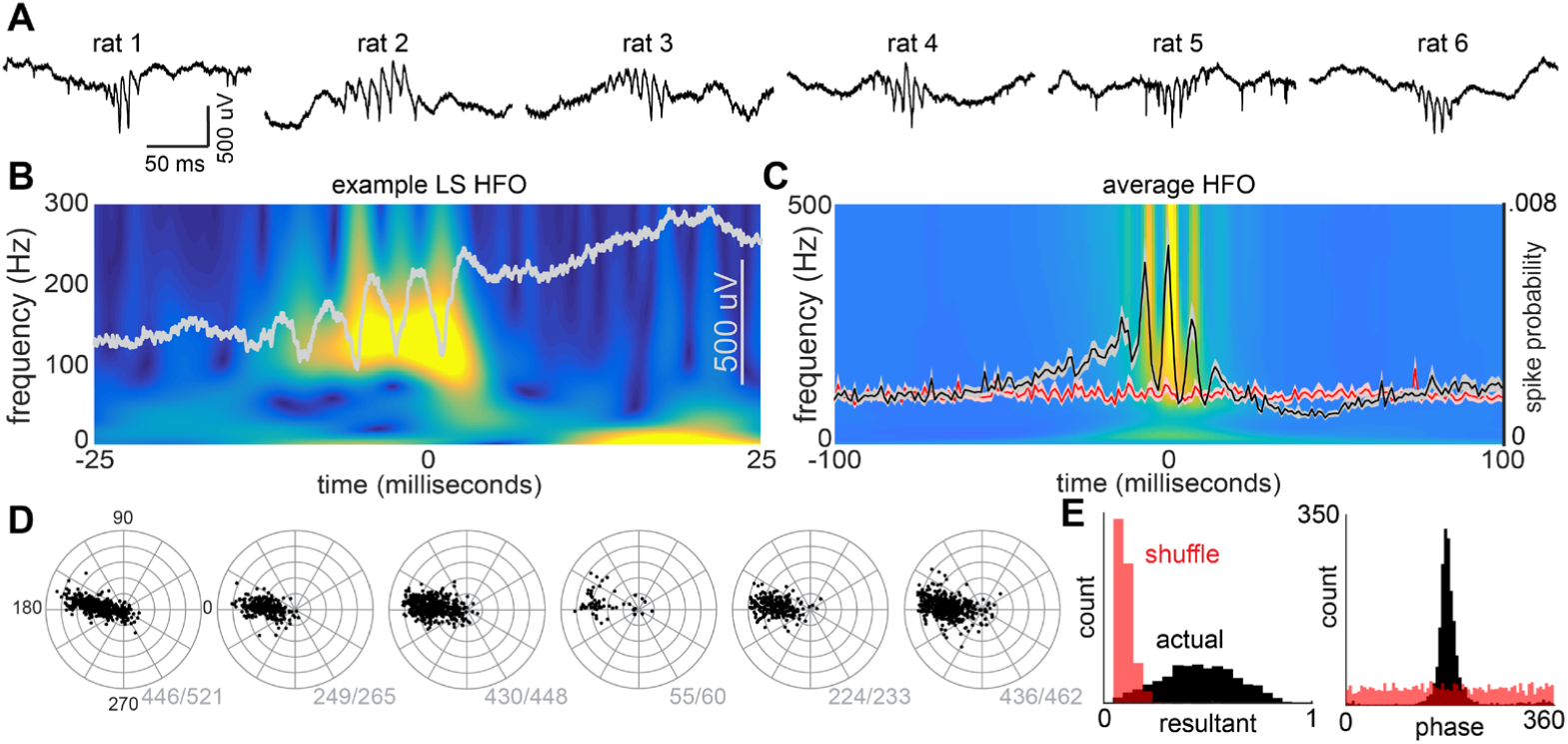
HFO events in LS are synchronized population bursts. **(A)** Example LS high frequency oscillations (HFOs). Each raw trace is an example event from one animal (20 kHz). **(B)** Example power spectrum (wavelet transform) for a single LS HFO event (grey overlay is the raw trace). **(C)** Average power spectrum and LFP across all events (N = 6 animals, 29,955 events). *Overlay:* LS HFO event triggered LS spike probability, normalized by number of events (N = 1,989 neurons). Red line is the average spiking probability for an equivalent number of randomly selected segments for each recording. Bounds are +/- 3 SEM. **(D)** For each animal (N = 6), each black dot indicates the resultant magnitude and phase angle preference for a single neuron. 1840 of 1989 neurons were significantly (p < 0.05) phase-locked to 120-180 Hz oscillations. Radial axis is 0 to 1 (resultant length). Grey values are the number of significantly phase-locked neurons for each animal. **(E)** *Left:* Histogram of all resultant magnitudes (black) and shuffled control (red). *Right:* Histogram of mean phase angles (black) and shuffled control (red) for significantly phase locked neurons (P < 0.05).

We only observed HFO events at recording sites that were confined to the LS (Figure 2A-C). Across recordings in each animal, we adjusted the electrode sites to record from progressively deeper regions within LS (Figure 2D). Both the power of HFOs and the proportion of phase-locked neurons were highest in the intermediate LS and tapered off as the recordings approached either the dorsal or ventral boarder of LS (Figure 2E, F; S1). We also observed consistent temporal lags in LS HFOs between recording sites, suggesting a stereotyped direction of propagation common to all such events (Figure 2G-I). Across four animals where recording sites spanned the anterior-posterior axis, we observed temporal offsets with the posterior recording sites leading more anterior recording sites (Figure 2H). Temporal offsets could also be observed on electrode pairs aligned to the medial-lateral axis and the dorsal-ventral axis (Figure 2I). With these temporal offsets and the known distances between each pair of recording sites, we estimated that the travel speed was 1.01 meters/second in the posterior-to-anterior direction (N = 4 animals), 0.53 meters/second in the medial-to-lateral direction (N = 1 animal), and 2.19 meters/second in the dorsal-to-ventral direction (N = 4 animals). Together, these data suggest that the LS HFO is a travelling wave that propagates along a vector that largely aligns with the hippocampal afferent projections into the LS and efferent projections towards hypothalamic regions (Risold and Swanson, 1997; Swanson and Cowan, 1979; Tingley and Buzsáki, 2018).

**Figure 2.**
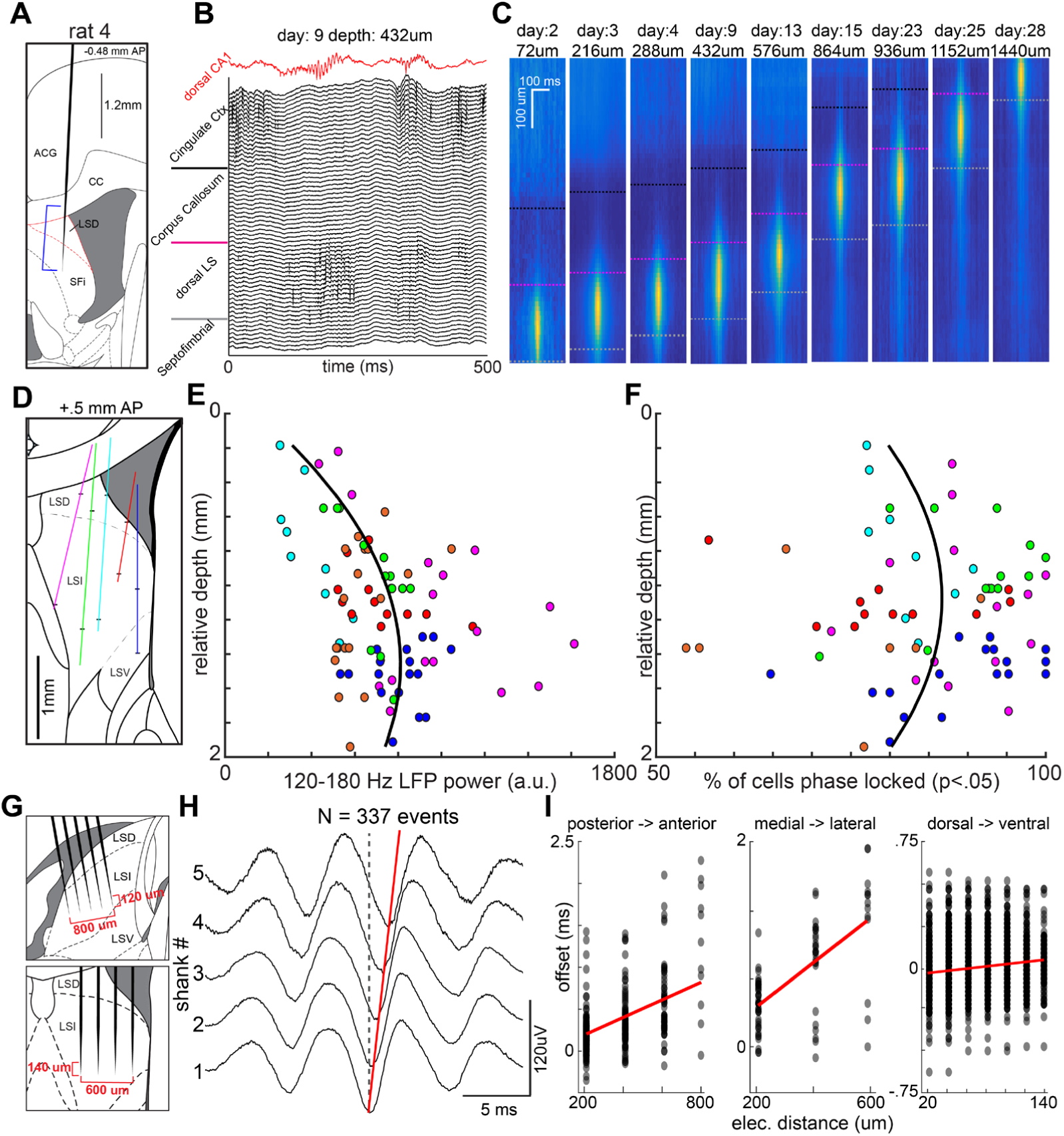
Localization and directionality of LS HFOs. **(A)** Single-shank silicon probe placement for one animal. Blue bracket indicates the span of 64 recording sites, spaced over 1260 µm for a single day. **(B)** Raw LFP traces for all 64 channels during an LS HFO event spanning multiple brain regions. Red trace is simultaneously recorded LFP from CA1 pyramidal layer. **(C)** Heat maps of 120-180 Hz power during LS HFO’s across days and recording depths. Black line, border between corpus callosum and LS; pink and white interrupted lines, span of recording sites in LS. **(D)** Diagram of recording tracks from 5 animals. **(E)** 120-180 Hz power of LS HFO events for each animal and recording depth. **(F)** The proportion of neurons significantly phase-locked (P < 0.05) to 120-180 Hz oscillations for each animal and recording depth. Colors for each dot match the respective tracts in panel D. Black lines are the best polynomial fit (order = 2). **(G)** *Top:* Sagittal view of lateral septum, and silicon probe layout spanning AP and DV axes. *Bottom:* Coronal view of the lateral septum, and silicon probe layout spanning ML and DV axes. **(H)** Example average LS HFO across 5 shanks oriented along the AP axis (N=337 events, 200 um spacing). Red line follows the trough across temporal delays. **(I)** For each pair of recording electrodes aligned to one axis (AP, ML, or DV; 20-800 µm apart), the temporal lag between troughs was calculated (y-axis) and compared with the physical distance between electrodes (x-axis). *Left:* LS HFOs are travelling waves in the posterior-to-anterior direction. *Middle:* LS HFOs are travelling waves in the medial-to-lateral direction. *Right:* LS HFOs are travelling waves in the dorsal-to-ventral direction.

Due to the alignment of these travelling waves with the innervation pattern of hippocampal fibers, and the fact that the hippocampus provides the main excitatory drive to LS (Risold and Swanson, 1997; Tingley and Buzsáki, 2018), we hypothesized that these events may be coupled in some way with hippocampal activity. Across recordings with sleep and behavioral sessions, we primarily observed HFO events during NREM sleep (Figure 3A). While this is similar to the state dependent occurrence of hippocampal SPW-Rs (Buzsáki et al., 1983), the overall rates of LS HFOs were significantly lower and the event durations were significantly shorter than hippocampal SPWRs (Figure 3B, C). Next, we examined the finer timescale relationship between hippocampal SPW-Rs and LS HFOs (Figure 3D). While some events were observed to occur independently (Figure 3D; left two examples), many LS HFOs occurred in close temporal proximity to dorsal hippocampal SPW-Rs (Figure 3D; right two examples). Across recordings, we observed a reliable coupling of events across structures (Figure 3E). Additionally, the strength of coupling between hippocampal SPW-Rs and LS HFOs was dependent upon the brain state of the animals. HPC-LS coupling was significantly stronger during NREM sleep than during the waking immobile state (two-way T-test; P<.01; Figure 3E, F). This state-dependent coupling could also be observed when examining the percentage of LS HFO events that occurred within ±25 milliseconds of a CA1 ripple (NREM = 0.3; WAKE = 0.15; P < 6.6e-6 two-way T-test; Figure 3F) and in the SPW-R triggered LS firing rates (Figure 3G).

**Figure 3.**
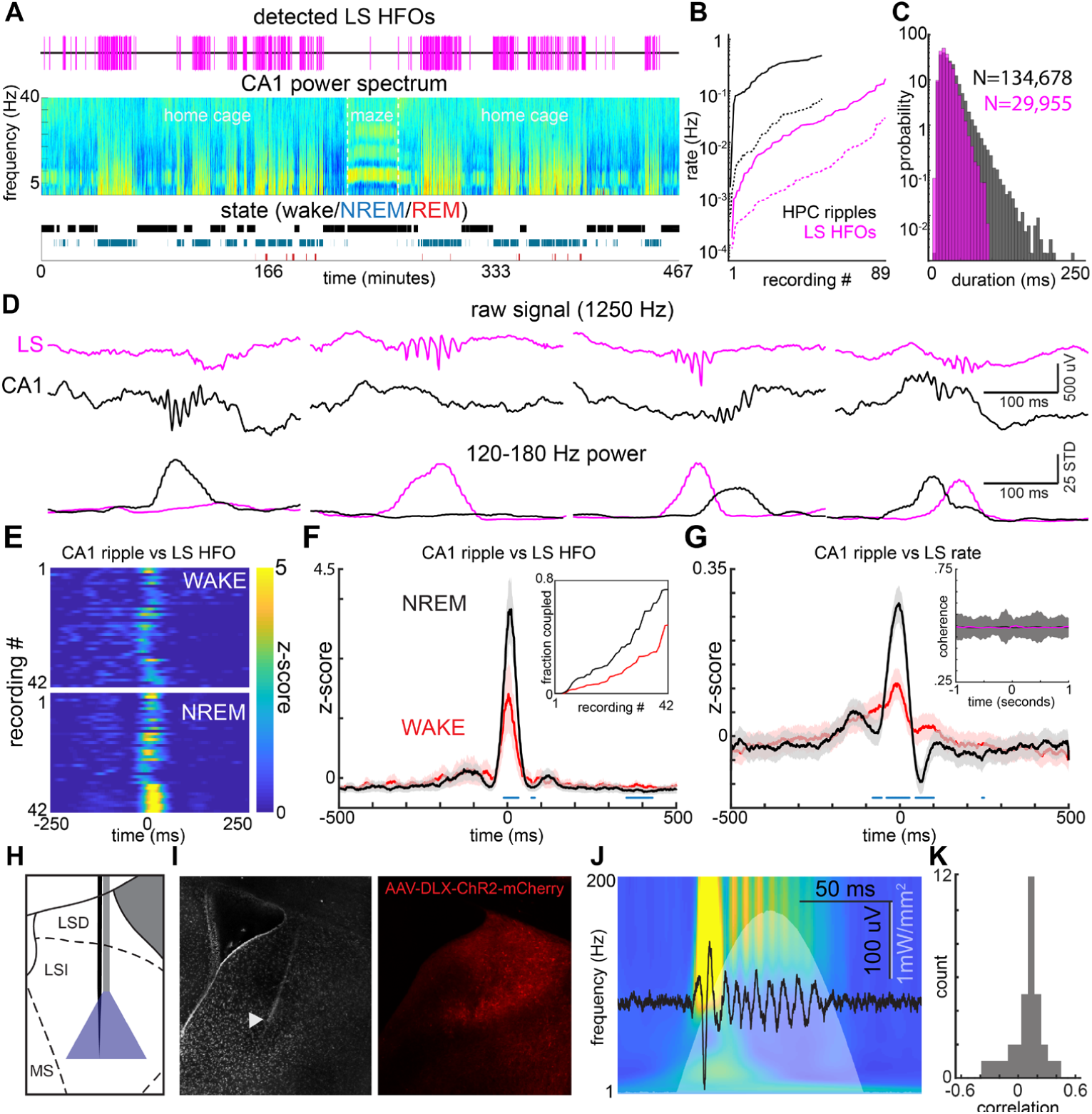
State dependence, coupling, and production of LS HFOs. **(A)** LS HFO occur more often during NREM sleep. *Top:* Raster of LS HFO occurrences (magenta lines). *Middle:* CA1 Power spectrum across ∼7.5 hour recording and the associated state scoring. *Bottom*: Scored state for the duration of the recording. **(B)** LS HFOs occur primarily during SWS. For each recording, the rate of LS HFOs (magenta; N = 89 recordings) during wake (dashed) and NREM (solid) are shown. Same for CA1 ripples in black (N = 53 recordings). **(C)** Duration histograms for CA1 ripples (black) and LS HFOs (magenta) across all detected events. **(D)** Four example events from simultaneous CA1-LS recordings. *Top:* Raw traces for LS (magenta) and CA1 (black). *Bottom:* 120-180 Hz Z-scored power for LS and CA1 channels. **(E)** Cross-correlogram of CA1 ripples with LS HFO events during wake (top panel) and NREM sleep (bottom panel) for all recordings. **(F)** Average cross-correlogram between CA1 ripples and LS HFOs (N = 42 recordings). Blue dots indicate bins with significantly different coupling during NREM sleep (two-way T-test; p < 0.001) *Inset:* Coupling of CA1 ripples and LS HFOs (within ±25 ms) is significantly higher during NREM sleep. **(G)** Average cross-correlogram between CA1 ripples and LS neuron firing rates (N = 1,313 neurons). Black trace is CA1 ripples during NREM sleep, red trace is for CA1 ripples during wake. *Inset:* Magenta is CA1-LS average 120-180 Hz phase coherence relative to CA1 ripples. Black trace is an equivalent number of randomly jittered events. Bounds are ±3 standard deviations. **(H)** Experimental paradigm for optogenetic experiments **(I)** Histological verification of recording site (left) and virus expression (right). **(J)** Black trace is one example LS response to a 100 millisecond Gaussian light pulse (white trace). Heat map is the average wavelet transform across 121 identical stimulations (rows are Z-scored). Color axis is −1 to 4 **(K)** For each stimulation protocol that elicited a response, the correlation between the power spectrum slope and the magnitude of each response is shown (N = 35).

There are at least two potential underlying mechanisms to explain the presence of HFOs in the lateral septum given the current data. First, LS HFOs may be ‘inherited’ from the hippocampus, such that the ripple oscillation itself is transferred to the LS wave-by-wave. Alternatively, the spiking output from the hippocampus, during SPW-Rs, provides the necessary level of excitatory drive for the induction of a locally generated oscillation in LS. Three observations argue in favor of local origin of LS HFOs. First, LS HFOs were significantly shorter than hippocampal SPW-Rs (Figure 3C) and were accompanied by a suppression of firing immediately following the population burst (Figures 1C, 3G). A direct inheritance of the hippocampal SPW-R would predict similar duration and firing patterns. Second, LS spiking activity was not phase-locked to individual waves of SPW-Rs and LS LFPs and phase coherence between HFOs and SPW-Rs was not significantly different from a shuffled distribution (Figure 3G). Third, optogenetic activation of the LS GABAergic population induced HFOs, reminiscent of the spontaneous events (Figure 3H-J). Using a GABAergic specific promoter, we expressed ChR2 in a majority of the LS population (Figure 3I; (Dimidschstein et al., 2016)). Using a Gaussian waveform of blue light (450 nm), mimicking the envelope of SPW-R-induced hippocampal excitation, local oscillations of ∼160 Hz could be reliably induced (Figure 3J; S2). These LS responses were strongly state dependent, with larger magnitude responses occurring when the power spectrum slope was steeper (i.e. when the animal was in NREM sleep; Figure 3K).

These findings demonstrate that LS populations can integrate hippocampal SPW-R activity and transmit shorter duration but highly synchronized population bursts to its hypothalamic and brainstem targets. Therefore, we examined which features of the hippocampal SPW-R best predicted responses of LS neurons. The experimental paradigm and data (Figure 4A-C) allowed for the extraction of several candidate features for every hippocampal SPW-R (Figure 4D). The first feature we examined was the number of action potentials elicited by the hippocampal population during SPW-Rs. For many LS neurons (38%), this measure significantly predicted the number of action potentials elicited in the postsynaptic LS neuron (Figure 4D,E). The next feature we examined was the average anatomical depth of neurons participating in a SPW-R, within the CA1 cell body layer (i.e. deep to superficial). For a set of LS neurons (23%), the recording depth of the active CA1 population correlated significantly with the number of action potentials elicited (Figure 4D,E). The brain state of the animal (Wake/NREM) within which the SPW-R occurred was yet another factor. As expected from the data presented in Figure 3, the rate of spiking for some LS neurons (26%) was significantly predicted by brain state (Figure 4E). The duration of hippocampal events could also be used to significantly predict firing rates for ∼20% of the LS neurons (Figure 4E).

**Figure 4.**
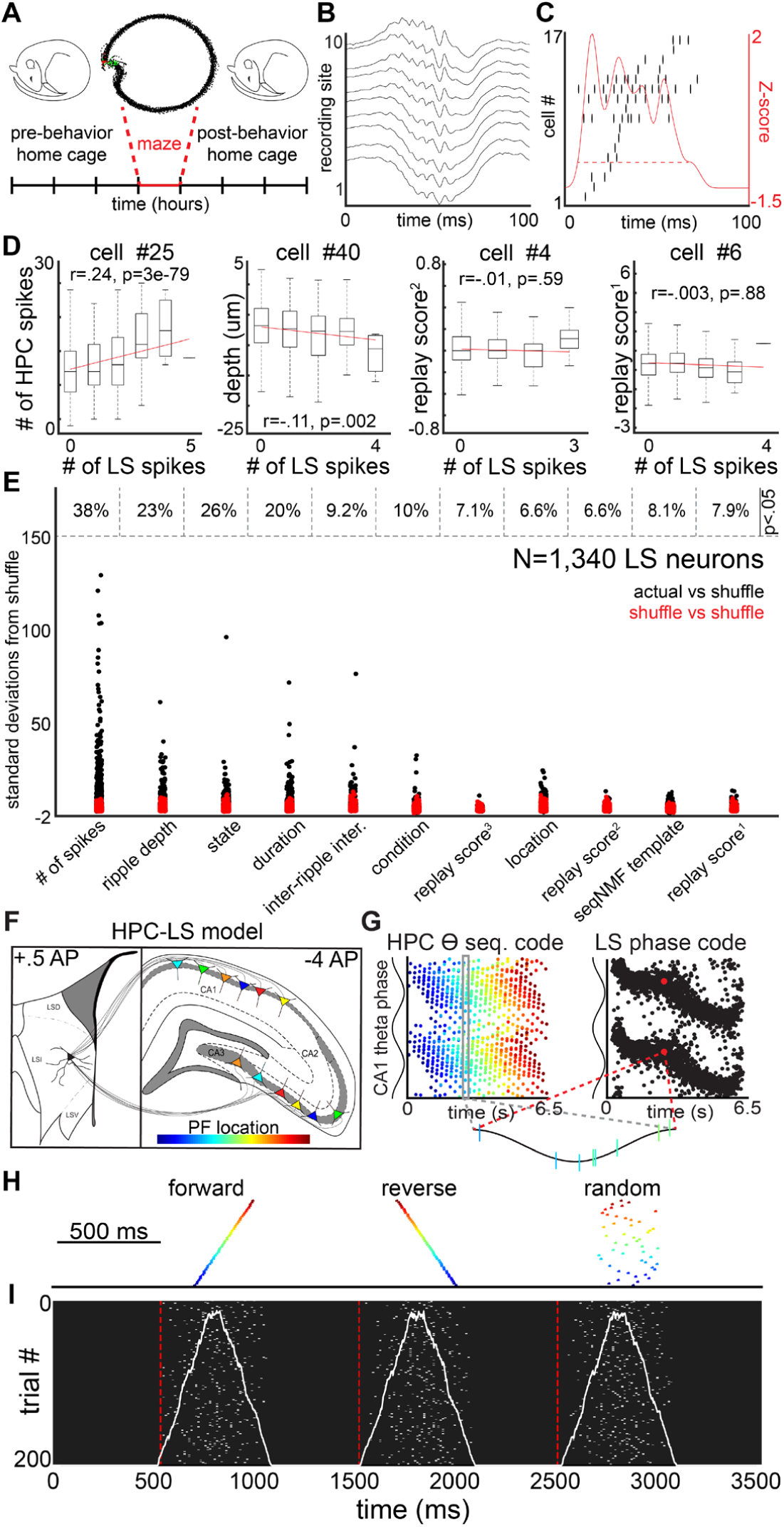
LS neuron firing is predicted by the magnitude and anatomical location of HPC SPWRs, but not ripple ‘content’. **(A)** Example diagram of experimental paradigm for all recordings. **(B)** LFP traces of a single HPC SPW-R. **(C)** Spiking activity from 17 CA1 neurons during the ripple shown in C and their summed (Z-scored) firing. **(D)** Four example LS neurons showing the relationship between LS firing (X-axis) and number of CA1 spikes, recording depth, and replay scores, respectively (Y-axes). Examples were selected from neurons within ± 5% of the mean for each distribution. **(E)** Ripple features examined to predict LS neuron firing (X-axis). Y-axis is the number of standard deviations that the mean squared error (MSE) was from a distribution of MSE values calculated from shuffled data for the same neuron. For every feature, each neuron is shown twice, as a black dot (actual data vs shuffled distribution) and as a red dot (one shuffled iteration vs shuffled distribution). The percentage of neurons surpassing p < 0.05 (two-way T-test) are shown above for each ripple feature. **(F)** Biophysical model of CA1/CA3 place field populations that provide convergent input to a LS neuron. Colors indicate the location of place fields along a simulated track. **(G)** The HPC-LS circuit transforms a population theta sequence code into a rate-independent phase code. **(H)** Forward, reverse, and scrambled ‘replay’ sequences were created in the presynaptic CA1/CA3 populations and the output LS neuron was examined. **(I)** Across trials with different jittered noise (Y-axis), the firing dynamics of the postsynaptic LS neuron (X-axis) were indistinguishable for different compressed spiking sequences

Neuronal sequences that occur during active experience are thought to be ‘replayed’ during hippocampal SPW-Rs (Skaggs and McNaughton, 1996). We, therefore, asked whether LS neurons can specifically ‘read out’ this precise temporally organized spiking. We used three replay quantification methods that have been previously developed: 1) the rank-order correlation between spiking order during each ripple with the firing rate maps from maze traversal (Diba and Buzsáki, 2007; Foster and Wilson, 2006)(replay score 1). 2) the maximized integral under the line of best fit (i.e. the discrete approximation of the Radon transform; (Toft, 1996)), of the posterior probability matrix calculated with a Bayesian approach (Davidson et al., 2009)(replay score 2), and 3) the linear weighted correlation of the posterior probability matrix calculated with a Bayesian approach (Grosmark and Buzsáki, 2016)(replay score 3). The firing rates of only a minority of LS neurons—7.9, 6.6, and 7.1% respectively—could be significantly predicted by the sequential structure of hippocampal events captured by these methods (Figure 4D,E).

As the proportion of hippocampal SPW-Rs that can be decoded as ‘replay’ events is small in the hippocampus (8-20%; Davidson; other REFS), we reasoned that perhaps some independent sequential structure in other hippocampal SPW-Rs may drive LS neuron firing. To examine this possibility, we took an unsupervised variance decomposition approach (Mackevicius et al., 2019). This method allowed us to extract sequential activity patterns from hippocampal SPW-Rs without relying on a pre-existing template. Across recordings, we were able to capture >95% of the variance in hippocampal SPW-R spiking activity with 40 ‘template sequences’ (Figure S3). We could then ask whether any of these identified templates correlated with firing of LS neurons. As with the ‘replay’ scores, we found that that the percentage of LS neurons whose action potential firing could be significantly predicted with these templates was small (Figure 4E; 8.3%; P<0.05).

We also observed a systematic increase in firing rates during post-behavior NREM sleep in CA1, CA3, and LS populations (Figure S4A, B). We did not observe fast timescale reactivation of hippocampal-LS or LS-LS cell pairs that exceeded what would be expected by chance, given the observed changes in firing rate (Figure S4C).

Thus, responses of LS neurons to hippocampal drive is related most strongly to the magnitude of synchronized excitation (i.e., the number of spikes within a SPW-R), rather than the precise temporal organization within particular SPW-R events. To further examine this phenomenon we utilized a model of the hippocampus-LS circuit that replicates previously described in vivo rate-independent phase encoding of spatial position ((Tingley and Buzsáki, 2018); Figure 4F, G). During simulated ‘maze traversals’ this model is capable of reading out hippocampal theta sequences and converting them into a pure spike-phase, rather than spike-rate, code for position (Figure 4G). By providing the model with temporally compressed forward, reverse, or scrambled replay events (Figure 4H), we observed that the response of the postsynaptic model LS neuron was nearly identical for all three event types (Figure 4I). Thus, during periods of high excitatory drive, information about the sequential ordering of hippocampal neurons is lost and the LS neurons’ firing rate best corresponds to the magnitude of this depolarization.

## Discussion

We demonstrated the presence of a locally generated high frequency oscillation in the LS. Many, but not all, LS HFOs co-occurred with dorsal hippocampal SPW-Rs with the highest probability of coupling occurring during NREM sleep. This appears to be the inverse relationship of what is observed between hippocampus and prefrontal cortex (Tang et al., 2017), suggesting there is a state dependent routing of SPW-Rs to cortical or subcortical structures. It remains to be seen whether the ‘uncoupled’ events were in fact coupled with unmeasured intermediate or ventral hippocampal SPW-Rs, or some other unidentified brain region. The HFOs were localized exclusively to LS recording sites. Recordings more dorsal, in deep layers of anterior cingulate cortex, showed down state-coupled activity approximately 50 milliseconds after the CA1 SPW-R as has been previously observed (Sirota et al., 2003b). Recordings more medial/ventral, in medial septum, did not have such HFOs and showed primarily a suppression in firing rates, also in line with previous work (Dragoi et al., 1999). We also found that LS HFOs are traveling waves whose directionality closely matches the anatomical inputs coming from the hippocampus and its own outputs to hypothalamic and brainstem nuclei.

LS HFOs appear to have a different mechanism of generation than hippocampal or neocortical ripple oscillations. Our optogenetic experiments and the lack of phase coherence between structures suggest that a depolarizing envelope is all that is required for the local production of HFOs, while the ripple component of the SPW-R does not appear to be necessary for production. Thus, the high degree of convergence in this circuit, and the high level of synchronous spiking output during SPW-Rs provides the necessary level of excitation to locally induce these HFOs, without transferring the ripple oscillation in a wave-by-wave manner.

Finally, the ability of LS neurons to ‘read out’ hippocampal activity patterns depended mainly on the magnitude of synchrony of hippocampal pyramidal neurons during SPW-Rs, irrespective of their precise temporal organization. This can be contrasted with the theta state, where LS neurons are capable of reading out hippocampal sequences (Tingley and Buzsáki, 2018). These data and our modeling work here suggest that this difference can be explained by the greatly increased level of excitation that occurs during temporally compressed ‘replay’. Thus, the same LS circuit can (during theta) or cannot (during SPW-Rs) read out neuronal sequences, depending on the magnitude of neuronal synchrony.

Sequential activity during hippocampal SPW-Rs is hypothesized to be critical for consolidating past experiences and planning/imagining future action (Buzsáki, 2015; Diba and Buzsáki, 2007; Foster, 2017b; Pfeiffer, 2018). Within this framework, the neuronal sequences in the hippocampus call up and order neocortical cell assemblies. A requisite prediction of such a model is that postsynaptic neurons receiving unique hippocampal activity patterns must be able to differentiate events based on the temporal organization within spike sequences. Operating in parallel to this corticopetal mechanism, excitatory hippocampal output also massively converges on the neurons of the LS (Swanson and Cowan, 1975; Swanson et al., 1981) and, in turn, is conveyed to the motivation and locomotion organizing lateral hypothalamus and mesencephalon, respectively. Our findings demonstrate that this latter route is highly active during SPW-Rs. In turn, this subcortical funneling of hippocampal output, active primarily during consummatory behaviors and NREM sleep, can drive activity in regions that are known to modulate the metabolic and motivational state of animals through the endocrine (Del Rey et al., 2008) and peripheral nervous systems (Shimazu, 1987). The significance of these SPW-R-induced physiological effects remains to be explored.

## Acknowledgements

We would like to thank Antonio Fernández-Ruiz, Kathryn McClain, Daniel Levenstein, Azahara Oliva and Ipshita Zutshi for helpful comments on the manuscript.

## Supplemental Figures

**Figure S1.**
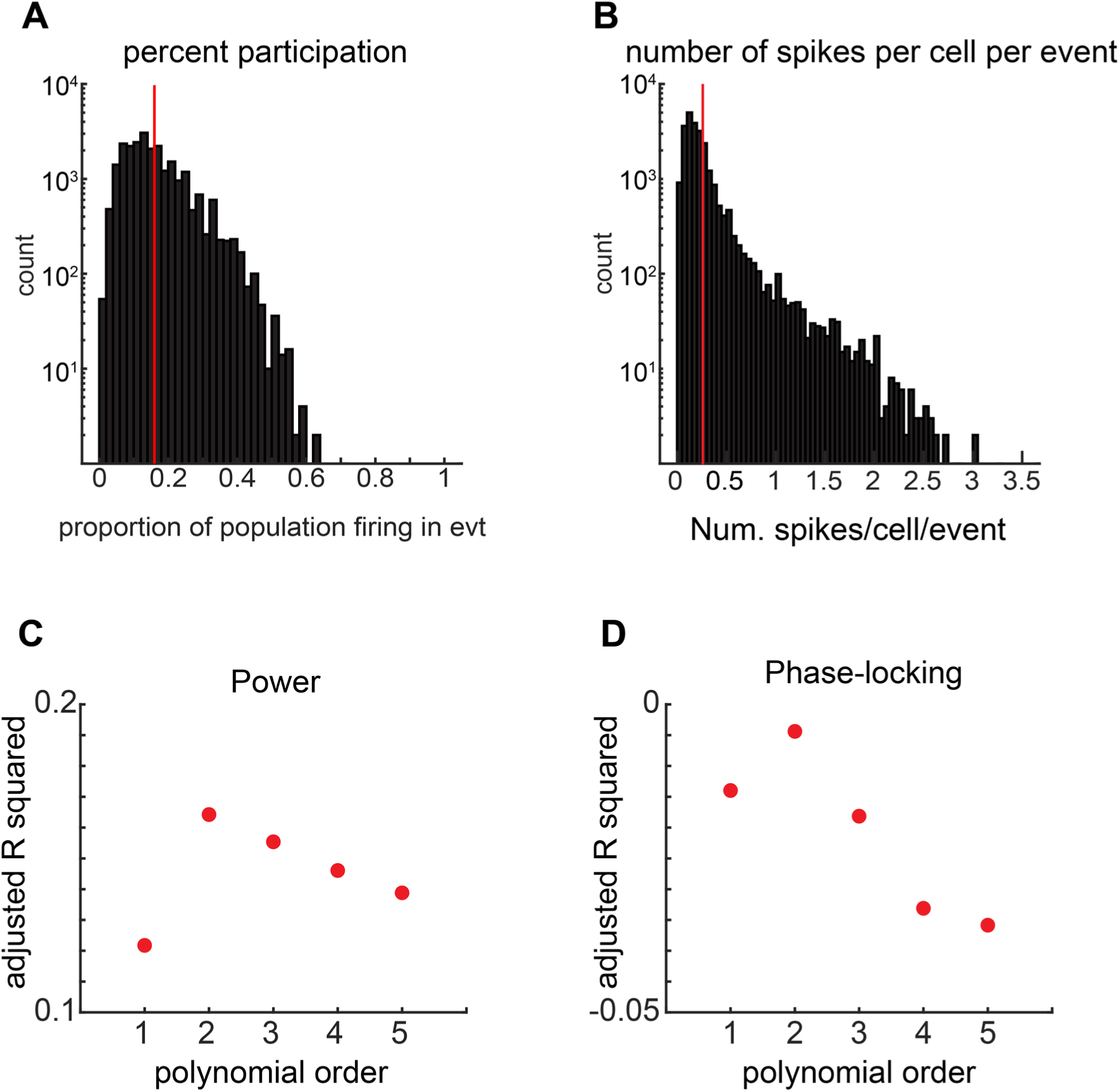
**(A)** Adjusted R-squared values for polynomial fits (order = 1-5) to the data in Figure 2E. The data are best fit by a polynomial of order 2. **(B)** Adjusted R-squared values for polynomial fits (order = 1-5) to the data in Figure 2F. The data are best fit by a polynomial of order 2.

**Figure S2.**
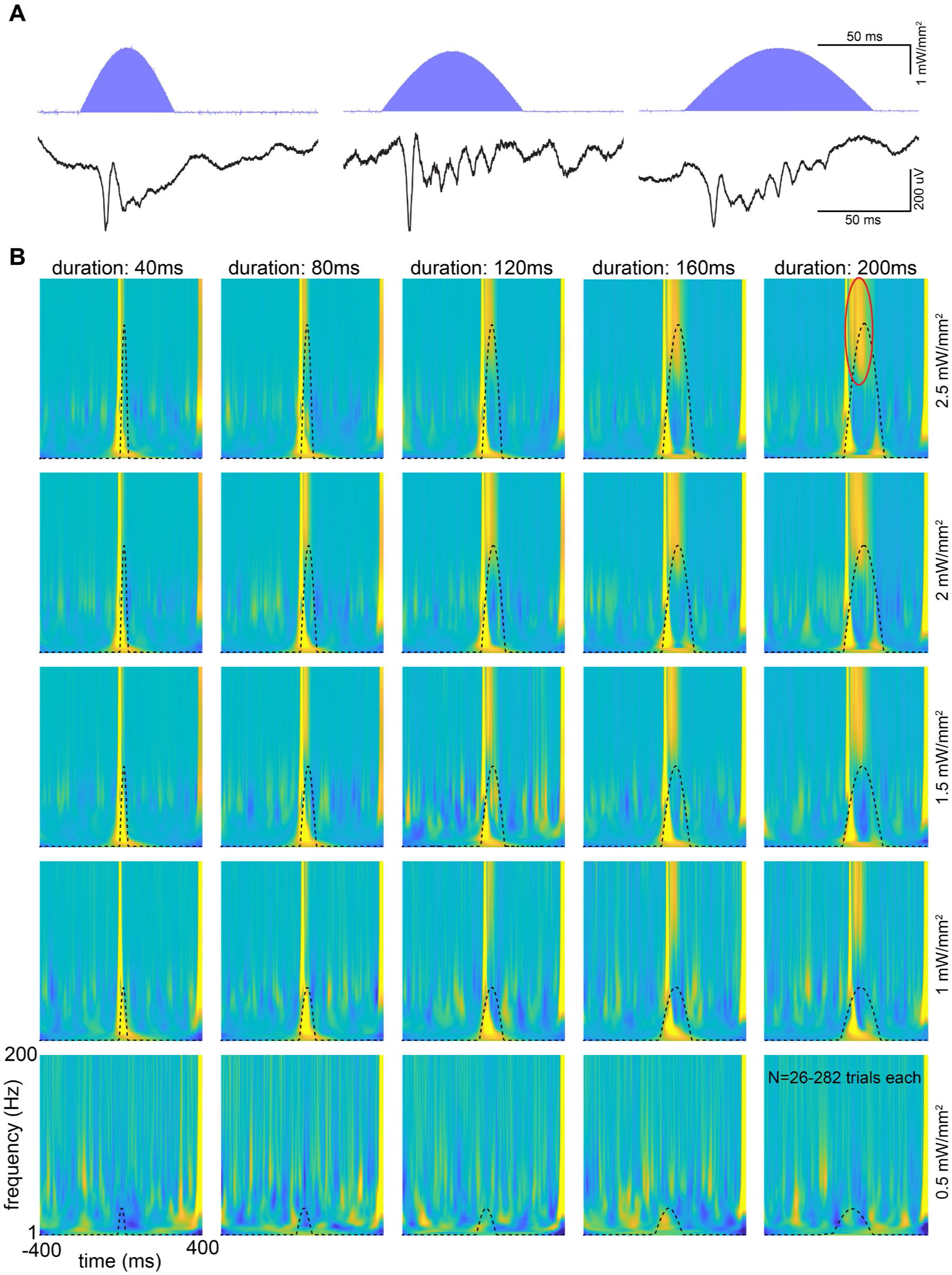
Optogenetic induction of LS HFOs (A) Black traces are example LFP responses to Gaussian light pulses of different durations (blue traces trace). **(B)** Duration and amplitude of stimulation were varied (40-200 milliseconds; 0.5-2.5 mW/mm^2^) to examine LS responses. Each heat map is the average wavelet transform of 26-282 stimulations of a given duration and amplitude. Each row is Z-scored, color axis is −4 to 4.

**Figure S3.**
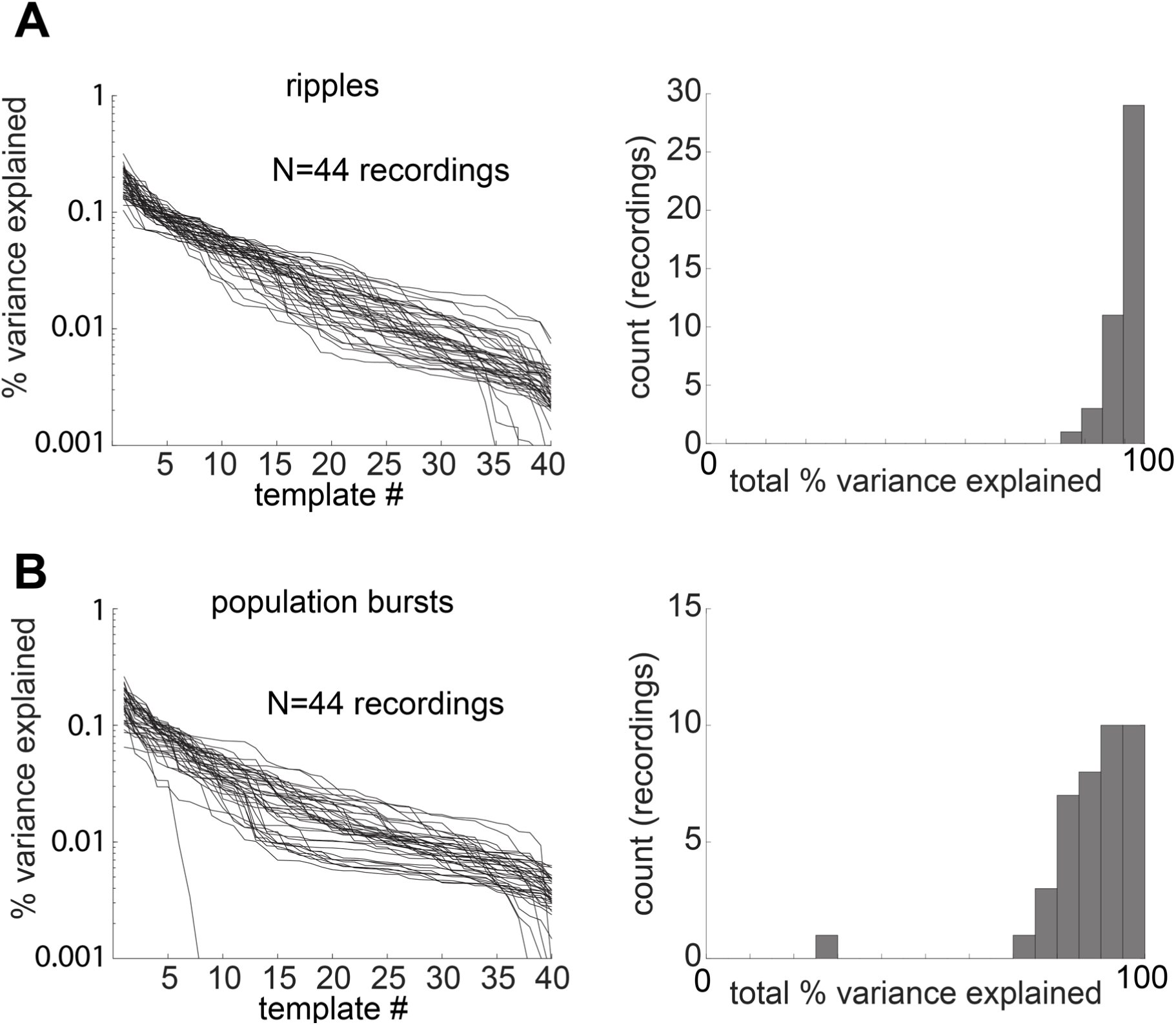
**(A)** Left: For all recordings (individual lines), each template derived from ripple events (X-axis) and the percent variance it explained from the data (Y-axis) is shown. Right: Histogram of the total percent variance explained across all 40 templates. Mean is 96%. **(B)** Left: For all recordings (individual lines), each template derived from population burst events (X-axis) and the percent variance it explained from the data (Y-axis) is shown. Right: Histogram of the total percent variance explained across all 40 templates. Mean is 88%.

**Figure S4.**
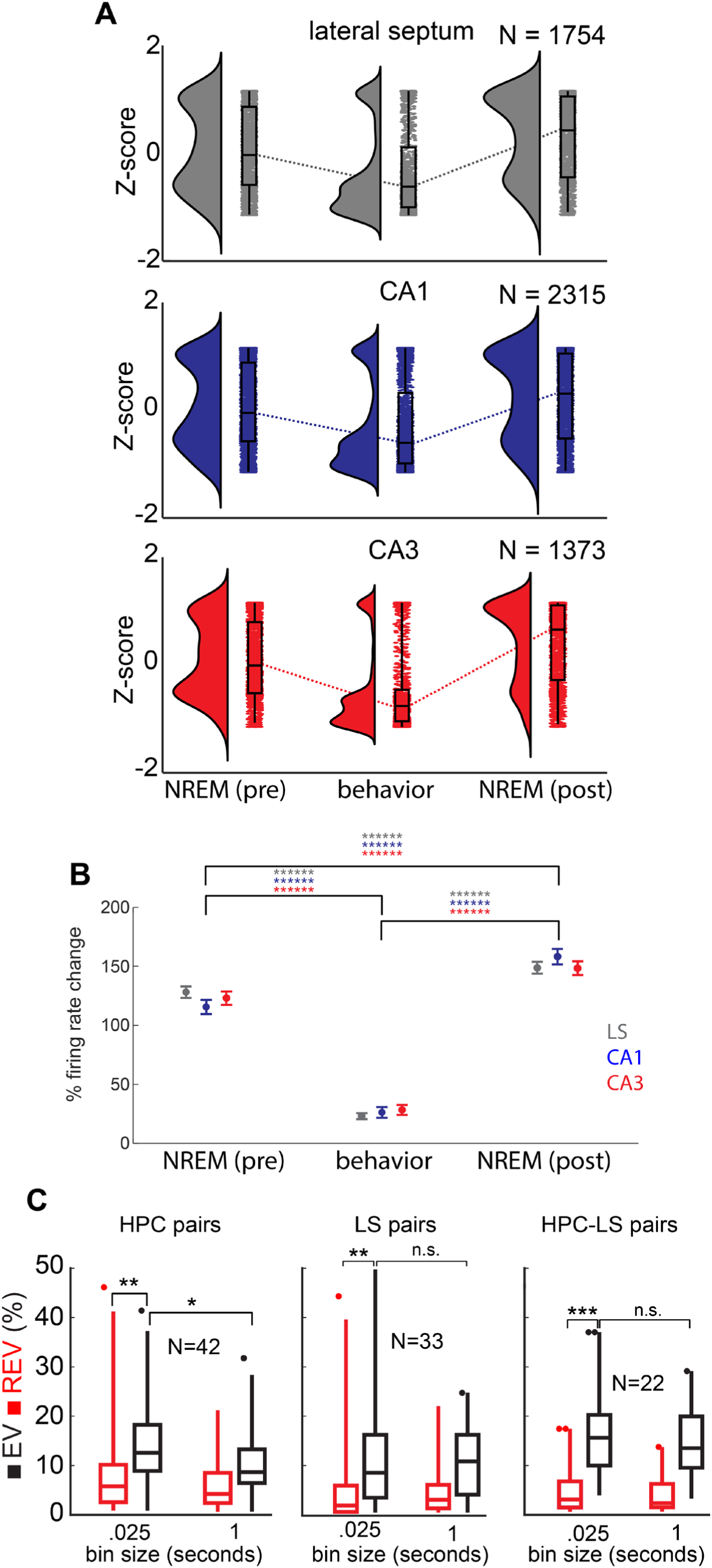
Systematic firing rate changes, but no reactivation in LS. **(A)** Z-scored firing rates during NREM prior to behavioral session (left column), behavior (maze traversal; middle column), and NREM after behavioral session (right column) for lateral septum (top), CA1 (middle), and CA3 (bottom). **(B)** Average changes in firing rate (relative to whole session average) during NREM prior to behavioral session (left column), behavior (middle column), and NREM after behavioral session (right column) for lateral septum (green), CA1 (red), and CA3 (blue). Bounds are ± three SEM. **(C)** For all pairs of neurons (Left: HPC-HPC; Middle: LS-LS; Right: HPC-LS) the explained variance and reverse explained variance are shown for .025 second bins and 1 second bins. All * signify P < 0.05 using two-way T-tests.

## SUPPLEMENTARY METHODS

The experiments were performed on the same Long-Evans rats 3 males and 2 females) which were described in Tingley and Buzsaki (2018). Detailed description of behavioral training, surgical procedures and histological results are available there. Each recording consisted of three periods of time recorded continuously; 1) pre-behavior home cage, 2) maze traversal, and 3) post-behavior home cage. One additional animal, included here, was implanted with a tetrode Microdrive (rat 1) and included in the phase locking analysis. The dataset used for Figure 4 consists of 62 recordings with both CA1 and LS simultaneously recorded (N = 37) or CA3 and LS simultaneously recorded (N = 25). For CA3 recordings, a reference electrode placed in CA1 was used to detect SPW-Rs. No significant differences were observed when using CA1 or CA3 populations and these recordings were therefore merged. Ripple depth was not analyzed for CA3 populations.

### Recording/Data processing

Recordings were conducted using the Intan RHD2000 interface board, sampled at 20 kHz. Amplification and digitization were done on the head stage. Waveform extraction and initial clustering was conducted using SpikeDetekt and Klustakwik. Example parameters for these algorithms can be found in the GitHub repository (https://github.com/DavidTingley/papers). Manual waveform discrimination was then conducted using the Klustaviewa software suite. Waveform amplitude was utilized during this stage to assess unit stability. Any waveforms that changed significantly throughout the duration of the recording were discarded. Waveform isolation quality was quantified using the isolation distance metric (https://github.com/buzsakilab/buzcode) and the waveform amplitude.

For one animal, position within the environment was tracked with two head mounted LED’s (1 blue, 1 red) and an overhead camera (Basler, 30 Hz). For the other four animals, position was tracked with the OptiTrack camera system. IR reflective markers were mounted in unique positions on each animals’ head stage and imaged simultaneously by six cameras (Flex 3) placed above the behavioral apparatus. Calibration across cameras allowed for the three dimensional reconstruction of the animals’ head position, and head orientation, to within 1 mm (avg. displacement error = .70 mm ± 1.5 mm) at 120 Hz.

Position data was analyzed and segmented using a custom Matlab software suite. Only ballistic trials, without stopping or deviation from the trained trajectory, were extracted for further analysis. These trials made up ∼90-95% of all trials attempted for any given recording.

### Ripple detection

A modified version of a previously reported detection algorithm was used for both hippocampus and LS HFO detection (Csicsvari et al., 1999). Input parameters for the algorithm are stored in each of the MATLAB data structures storing ripple events. The code is available at https://github.com/buzsakilab/buzcode. Ripples were detected using the normalized squared signal (NSS) by thresholding the baseline, merging neighboring events, thresholding the peaks, and discarding events with excessive duration. Thresholds are computed as multiples of the standard deviation of the NSS. The estimated EMG was also used as an additional exclusion criteria (Schomburg et al., 2014). The same detection algorithm was used for both LS and HPC.

### Phase locking analysis

Epochs of time where the 120-200 Hz power was continuously above two standard deviations from the mean were identified. The phase angles for each timestamp where an action potential occurred within these epochs was calculated using the Hilbert transform. For each neuron, a histogram of phase angles was calculated for all action potentials that occurred within these LFP power thresholded epochs. The circular mean and resultant vector were then calculated using these histograms for each neuron. For a null distribution, LFP phase angles were circularly shifted by a random offset and the phase angle histograms were then recalculated (10 iterations).

### State scoring

A previously described semi-automated sleep scoring algorithm was used (Watson et al., 2016). In some cases, manual curation of algorithm parameters or specific segments of recordings was conducted in order to best identify brain state.

### Spatiotemporal patterns of population activity

The methods for quantifying activity patterns across hippocampal neural populations were taken from a variety of previous works. The ‘ripple-depth’ was quantified as the average anatomical depth (i.e., deep or superficial) for all neurons firing at least one action potential in a given event We used a previously developed method for estimating the position of recording sites, relative to the CA1 cell body layer (Mizuseki et al., 2011). This method uses the ripple band power envelope to estimate the position of the layer relative to recording sites. Each cell body position relative to the recording sites was then estimated by interpolating the action potential amplitudes across recording sites and selecting the largest amplitude at 5um increments. Aligning these two coordinates allows for the estimation of each recorded neuron relative to the CA1 cell body layer.

The inter-ripple interval for each event was defined as the mean of the durations immediately preceding, and following, a given ripple. The ripple ‘condition’ was defined as the experimental epoch (pre-behavior, behavior, or post-behavior) within which the event occurred. The replay scores consisted of 1) the rank-order correlation between spike times during spatial traversal and spike times of a given event (Foster and Wilson, 2006); 2) the ‘replay score’, which is the integral under the line of best fit, using the radon transform of the posterior probability matrix (Davidson et al., 2009); and 3) the ‘sequence score’, a normalized linear weighted correlation of the posterior probability matrix (Grosmark and Buzsáki, 2016).

A recently developed unsupervised method of spatiotemporal quantification of neural populations was also used (Mackevicius et al., 2019). This method allows for the examination of consistently occurring neural sequences without relying on a behavioral ‘template’ to relate too. Thus, if commonly ‘replayed’ spike sequences in HPC populations reliably predicted the firing dynamics of LS neurons, but were not tied to the experimental paradigm we imposed, this method allowed for the identification and analysis of such events.

### Reactivation analysis

The reactivation analysis was conducted as in (Kudrimoti et al., 1999). Explained variance (EV) was calculated by comparing pre-behavior NREM ripple epochs with behavioral epochs, while reverse explained variance was calculated by comparing post-behavior NREM ripple epochs with behavioral epochs.

However, as pair-to-pair correlations have been shown to fluctuate with firing rate ((De La Rocha et al., 2007)), this method is susceptible to fluctuations in basal firing rates between conditions ((Pavlides and Winson, 1989); Figure S4 A, B). Thus, we reasoned that altering the bin size for this analysis would allow us to capture only the slow time scale firing rate fluctuations (large bin size), and both the fast timescale reactivation and slow firing rate fluctuations (small bin size). We used bin sizes of 25 milliseconds for fast timescale reactivation, and 1 second to capture slower timescale fluctuations in firing rates.

Hippocampal populations showed significantly higher EV at fast (25 ms) timescales, suggesting the presence of reactivation. LS populations however, showed significantly higher correlations at slower timescales, suggesting that the observed increase in EV is due to changes in basal firing rate rather than network coordinated reactivation (Figure S4).

### Optogenetic experiments

One rat was injected with AAV-DLX-ChR2-mCherry in the right lateral septum. Two injections at 4.2mm and 4.6mm DV of 250uL each were conducted. After four weeks a 200-µm optic fiber and 50-µm tungsten recording wire were implanted. Square pulses and Gaussian waveforms were generated with a PulsePal device (Sanworks; v2) and transmitted to a laser diode driver (Thorlabs; LDC205C). Coupling efficiency was estimated using a photometer (Thorlabs; PM100D) and all power measurements provided are the mW/mm^2^ at the fiber tip prior to implantation.

